# Identification of *Acinetobacter baumannii* loci for capsular polysaccharide (KL) and lipooligosaccharide outer core (OCL) synthesis in genome assemblies using curated reference databases compatible with Kaptive

**DOI:** 10.1101/869370

**Authors:** Kelly L. Wyres, Sarah M. Cahill, Kathryn E. Holt, Ruth M. Hall, Johanna J. Kenyon

## Abstract

Multiply antibiotic resistant *Acinetobacter baumannii* infections are a global public health concern and accurate tracking of the spread of specific lineages is needed. Variation in the composition and structure of capsular polysaccharide (CPS), a critical determinant of virulence and phage susceptibility, makes it an attractive epidemiological marker. The outer core (OC) of lipooligosaccharide also exhibits variation. To take better advantage of the untapped information available in whole genome sequences, we have created a curated reference database of the 92 publicly available gene clusters at the locus encoding proteins responsible for biosynthesis and export of CPS (K locus), and a second database for the 12 gene clusters at the locus for outer core biosynthesis (OC locus). Each entry has been assigned a unique KL or OCL number, and is fully annotated using a simple, transparent and standardised nomenclature. These databases are compatible with *Kaptive*, a tool for *in silico* typing of bacterial surface polysaccharide loci, and their utility was validated using a) >630 assembled *A. baumannii* draft genomes for which the KL and OCL regions had been previously typed manually, and b) 3386 *A. baumannii* genome assemblies downloaded from NCBI. Among the previously typed genomes, *Kaptive* was able to confidently assign KL and OCL types with 100% accuracy. Among the genomes retrieved from NCBI, *Kaptive* detected known KL and OCL in 87% and 90% of genomes, respectively indicating that the majority of common KL and OCL types are captured within the databases; 13 KL were not detected in any public genome assembly. The failure to assign a KL or OCL type may indicate incomplete or poor-quality genomes. However, further novel variants may remain to be documented. Combining outputs with multi-locus sequence typing (Institut Pasteur scheme) revealed multiple KL and OCL types in collections of a single sequence type (ST) representing each of the two predominant globally-distributed clones, ST1 of GC1 and ST2 of GC2, and in collections of other clones comprising >20 isolates each (ST10, ST25, and ST140), indicating extensive within-clone replacement of these loci. The databases are available at https://github.com/katholt/Kaptive and will be updated as further locus types become available.

**Data Summary:** 1. Databases including fully annotated gene cluster sequences for *A. baumannii* K loci and OC loci are available for download at https://github.com/katholt/Kaptive

2. The *Kaptive* software, which can be used to screen new genomes against the K and O locus database is available at https://github.com/katholt/Kaptive (command-line code) and http://kaptive.holtlab.net/ (interactive web service).

3. Details of the *Kaptive* search results validating *in silico* serotyping of K and O loci using our approach are provided as supplementary files, Dataset 1 (92 KL reference sequences and 12 OCL reference sequences), Dataset 2 (642 genomes assembled from reads available in NCBI SRA) and Dataset 3 (3415 genome assemblies downloaded from NCBI GenBank).

**Impact statement:** The ability to identify and track closely related isolates is key to understanding, and ultimately controlling, the spread of multiply antibiotic resistant *A. baumannii* causing difficult to treat infections, which are an urgent public health threat. Extensive variation in the KL and OCL gene clusters responsible for biosynthesis of capsule and the outer core of lipooligosaccharide, respectively, are potentially highly informative epidemiological markers. However, clear, well-documented identification of each variant and simple-to-use tools and procedures are needed to reliably identify them in genome sequence data. Here, we present curated databases compatible with the available web-based and command-line *Kaptive* tool to make KL and OCL typing readily accessible to assist epidemiological surveillance of this species. As many bacteriophage recognise specific properties of the capsule and attach to it, capsule typing is also important in assessing the potential of specific phage for therapy on a case by case basis.

## Introduction

One of the most imminent global health crises is the increasing prevalence and global dissemination of highly resistant bacterial pathogens that are able to persist in hospital environments despite infection control procedures. In 2017, the World Health Organisation identified carbapenem-resistant strains of the opportunistic Gram-negative bacterium, *Acinetobacter baumannii*, as a critical priority for therapeutics development due to alarming levels of resistance against nearly all clinically suitable antibiotics (1). The success of extensively antibiotic resistant *A. baumannii* isolates can be attributed, in part, to the evolution and expansion of well adapted clonal lineages (2-5), including the two major globally disseminated clones, Global Clone 1 (GC1) and Global Clone 2 (GC2), and other lineages that are found less frequently (e.g. sequence type 25; ST25) or on only one or two continents (e.g. ST78) (6). Hence, the development of precise epidemiological tracking methods for *A. baumannii* isolates, in particular those from important clonal lineages, are urgently needed to enhance surveillance and improve our understanding of how *A. baumannii* circulates both locally and globally.

Traditionally, epidemiological studies tracing important bacterial lineages associated with human and animal infections used serological typing of the polysaccharides produced on the cell surface (7, 8), as there can be significant variation in structures observed on different isolates of the same species (9-12). The cell-surface polysaccharides targeted in these schemes included capsular polysaccharide (CPS, K, or capsule) and/or O-antigen polysaccharide (OPS or O) that is attached to lipooligosaccharide (LOS) forming a lipopolysaccharide (LPS). In early studies, an *A. baumannii* serological typing scheme was developed for a major immunogenic polysaccharide, believed at the time to be the O antigen (13, 14), and 38 different serovars were included in the last update to the scheme nearly two decades ago (15). However, this system is no longer used.

In the last decade, it has been shown that the major immunogenic polysaccharide produced by the species is CPS not O antigen (16-18). The CPS of *A. baumannii* is a major virulence determinant as isolates lacking CPS do not cause infections (17). CPS is also a key target of potential novel control strategies including phage therapy (19, 20) and vaccinations (21, 22). Unfortunately, the current lack of knowledge about capsule diversity and epidemiology in the broader *A. baumannii* population, and lack of tools to readily detect changes in the population distribution hinders effective design of these controls.

Most of the genes that direct the synthesis of the CPS are clustered at the K locus (KL) that is located between the *fkpA* and *lldP* genes in the *A. baumannii* chromosome (16, 23). The general arrangement of the K locus features three main regions (Figure 1A). On one side, a module of genes for CPS export machinery (*wza-wzb-wzc*) are in a separate operon, divergently transcribed from the remainder of the gene cluster. On the other side lies a module of genes involved in the synthesis of simple sugar substrates. However, the *gne1* gene can be lost (e.g. Figure 2B) if D-Gal*p*NAc is not present in the CPS, and various other genes have been found between *gne* (or *gpi*) and *pgm* in some KL (24-26). The genetic content of the central region is specific to the CPS structure produced. It includes genes for the required number of glycosyltransferases, and the capsule processing genes (*wzx* and *wzy*). If complex sugars (e.g. pseudamininc acid, legionaminic acid, acinetaminic acid, bacillosamine, etc.) are included in the CPS, the central region will also contain genes for the synthesis and modification of these sugars (16, 27-30). Each distinct gene cluster, defined by a difference in gene content between *fkpA* and *lldP*, is assigned a unique identifying number (KL1, KL2, etc.). To date, more than 128 KL gene clusters (KL types) have been identified at the K locus in *A. baumannii* genomes (31).

**Figure 1.**
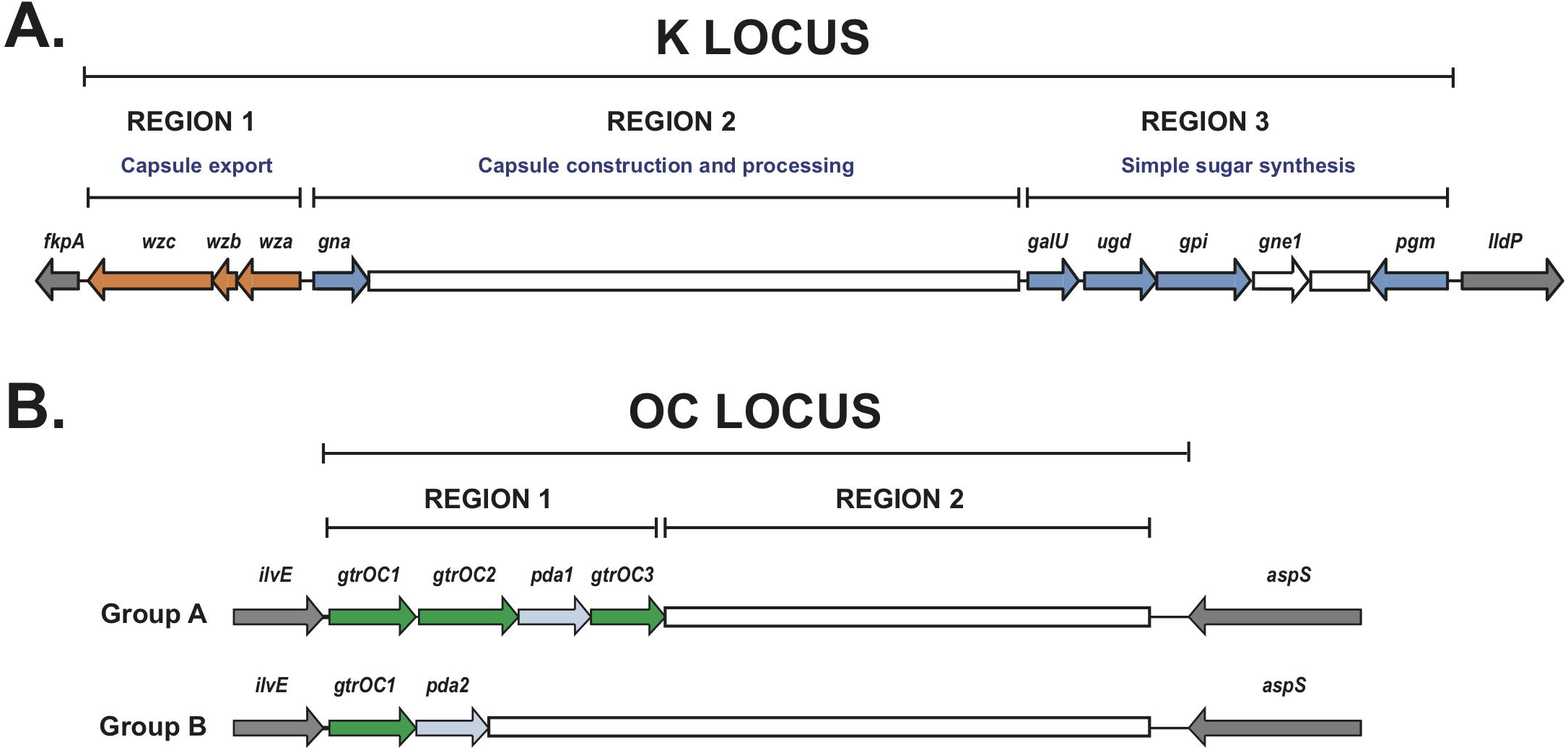
General arrangement of the surface polysaccharide synthesis loci in *A. baumannii.* KL and OCL boundaries are shown and flanking locus genes are coloured grey. Variable sequence portions are indicated by white boxes, and conserved genes at each locus are represented by coloured arrows. **A.** Organisation of the K locus with marked regions defining the roles of common modules. CPS export genes are orange and dark blue genes are involved in the synthesis of common sugar substrates. *gne1* is not always present but is often critical to the synthesis of many CPS structures. **B.** Organisation of the two groups (A and B) of the OC locus with marked regions defining conserved or variable portions. Green genes encode conserved glycosyltransferases and light blue are those involved in complex sugar synthesis

**Figure 2.**
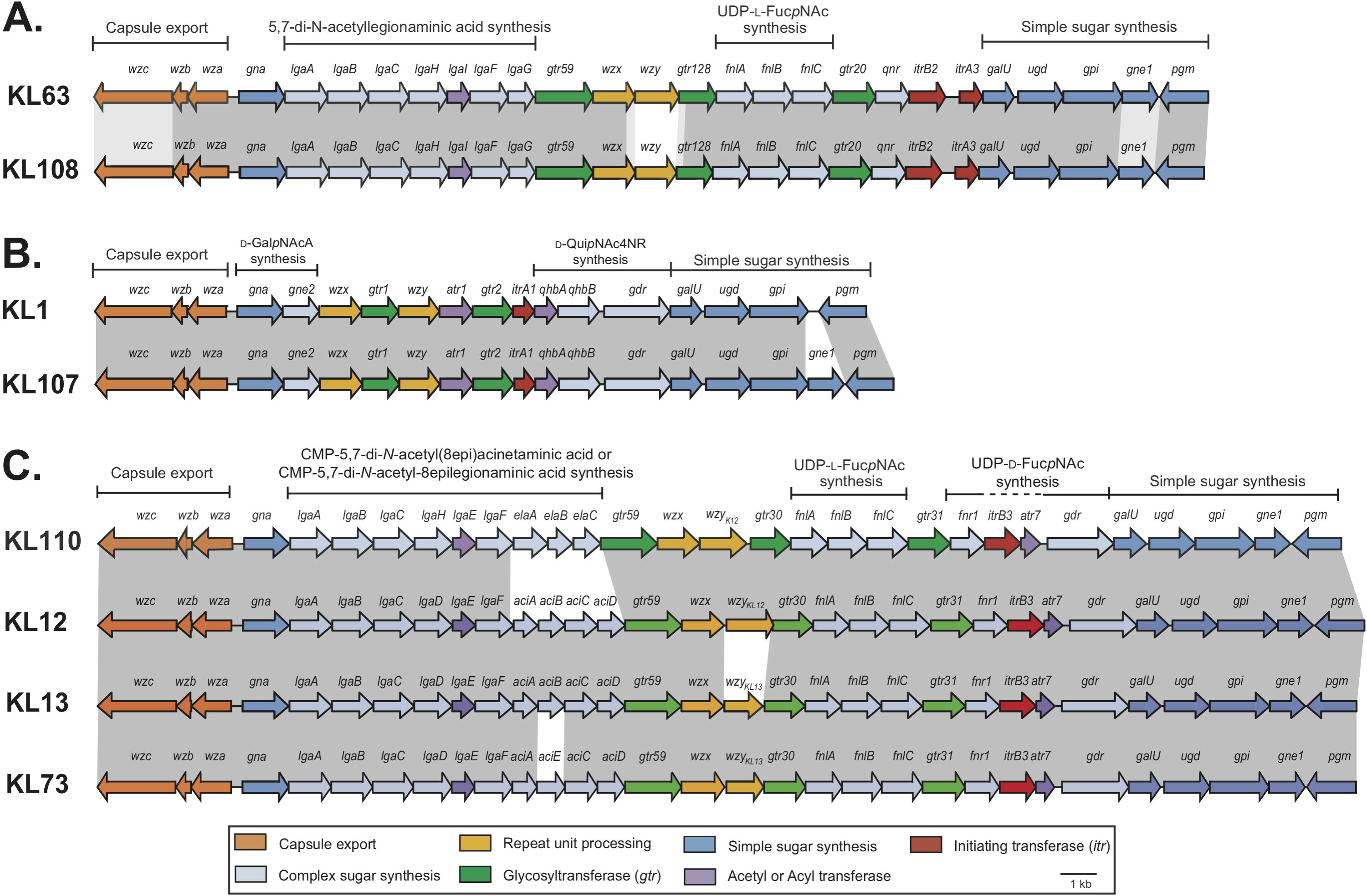
Closely related capsule biosynthesis gene clusters demonstrating cases of small genetic replacements. Genes are represented by arrows oriented in the direction of transcription that are coloured according to the scheme shown below. Shading between gene clusters indicates regions of >95% nucleotide sequence identity (dark grey) or 90-95% nucleotide sequence identity (light grey). Figure drawn to scale suing GenBank accession numbers listed in Table S1. **A.** KL63 and KL108 gene clusters differing in *wzy* sequence. **B.** KL1 and KL107 are an example of *gne1* presence vs. absence. **C.** KL13, KL73, KL12, and KL110 are examples of several closely related gene clusters with small sequence replacements altering the synthesis pathway of a complex sugar substrate, or topology of the CPS structure.

A transparent nomenclature system for CPS biosynthesis genes in *A. baumannii* was developed in 2013 to clearly identify the specific function of KL-encoded proteins for the non-expert (16). Where possible, gene names indicate enzyme function (i.e. Gtr assigned to <u>G</u>lycosyl<u>TR</u>ansferases and Itr to the transferases initiating K unit synthesis). For enzymes (e.g., Gtrs and Itrs) where sequence differences likely result in a change of substrate preference, a number indicating the different sequence type (cut off value of 85% aa sequence identity) is included in the name as a suffix. The current gene names are listed in Table 1. Most published annotations use this system (e.g. refs 26, 29-36). However, sometimes other nomenclature systems have been used (23, 37).

**Table 1.** Gene nomenclature key for *A. baumannii* K and OC loci

| Gene name | Predicted reaction product | Predicted protein |
| --- | --- | --- |
| **K locus** |  |  |
| *aci* | CMP-<u>A</u>cinetaminic <u>a</u>cid derivative | Multiple |
| *atr* | - | <u>A</u>cyl- or <u>A</u>cetyl- transferase |
| *alt* | - | D-<u>A</u>lanine transferase |
| *dga* | UDP-2,3-di<u>a</u>cetamido-2,3-dideoxy-D-glucuronic <u>a</u>cid | Multiple |
| *dmaA* | UDP-2,3-di<u>a</u>cetamido-2,3-dideoxy-D-mannuronic acid | 2-epimerase |
| *ela* | CMP-8-epi<u>l</u>egionaminic <u>a</u>cid derivative | Multiple |
| *fdt* | dTDP-D-Fuc<u>p</u>3NA<u>c</u> | Multiple |
| *fnl* | dTDP-L-Fuc<u>p</u>NA<u>c</u> | Multiple |
| *fnr* | UDP-D-Fuc<u>p</u>NA<u>c</u> | UDP-6-deoxy-4-keto-D-GalpNA<u>c</u> 4-<u>r</u>eductase |
| *galU* | UDP-D-Gl<u>c</u>p | UTP-glucose-1-phosphate uridylyltransferase |
| *gdr* | UDP-4-keto-6-deoxy-D-Gl<u>c</u>pNA<u>c</u> | UDP-Gl<u>c</u>pNA<u>c</u> 4,6-<u>d</u>ehydratase |
| *gna* | UDP-D-Gl<u>c</u>pNA<u>cA</u> | UDP-D-Gl<u>c</u>pNA<u>c</u> dehydrogenase |
| *gne1* | UDP-D-GalpNA<u>c</u> | UDP-D-Gl<u>c</u>pNA<u>c</u> epimerase |
| *gne2* | UDP-D-GalpNA<u>cA</u> | UDP-D-Gl<u>c</u>pNA<u>cA</u> epimerase |
| *gpi* | L-Fructose-6-phosphate | glucose-6-phosphate isomerase |
| *gtr* | - | Glycosyltransferase |
| *itr* | - | Initiating transferase |
| *lga* | CMP-<u>L</u>egionaminic <u>a</u>cid derivative | Multiple |
| *man* | GDP-D-mannose | Multiple |
| *mna* | UDP-D-Man<u>p</u>NA<u>c</u> | Multiple |
| *neu* | CMP-N-acetylneuraminic acid | Multiple |
| *pet* | - | Phosphoethanolamine transferase |
| *pgm* | D-Glucose-1-phosphate | Phosphoglucomutase |
| *pgt* | - | Phosphoglycerol transferase |
| *psa* | CMP-<u>P</u>seudaminic <u>a</u>cid derivative | Multiple |
| *ptr* | - | Pyruvyl transferase |
| *qdt* | dTDP-D-Quip3NA<u>c</u> | Multiple |
| *qhb* | UDP-D-QuipNA<u>c</u>4NH<u>b</u> | Multiple |
| *qnr* | UDP-D-QuipNA<u>c</u> | UDP-6-deoxy-4-keto-D-Gl<u>c</u>pNA<u>c</u> 4-<u>r</u>eductase |
| *rml* | dTDP-L-Rhamnose | Multiple |
| *tle* | dTDP-6-deoxy-L-<u>t</u>alose | dTDP-L-Rhamnose epimerase |
| *ugd* | UDP-D-Gl<u>c</u>pA | UDP-D-Gl<u>c</u>p dehydrogenase |
| *vio* | dTDP-4-acetamido-4,6-dideoxy-D-glucose | Multiple |
| *wza* | - | Outer membrane protein |
| *wzb* | - | Protein tyrosine phosphatase |
| *wzc* | - | Protein tyrosine kinase |
| *wzx* | - | Repeat unit translocase |
| *wzy* | - | Repeat unit polymerase |
| **OC locus** |  |  |
| *ahy* | - | Predicted <u>a</u>cyl<u>h</u>ydrolase |
| *gtrOC* | - | Glycosyltransferase (outer core) |
| *pda* | UDP-D-Gl<u>c</u>N | Polysaccharide <u>d</u>eacetylase |
| *ptrOC* | - | Pyruvyl transferase (outer core) |
| *wecB* | UDP-D-Man<u>p</u>NA<u>c</u> | UDP-D-Gl<u>c</u>pNA<u>c</u> C2 epimerase |

A second locus with variable gene content involved in the production of a surface polysaccharide (16) has been shown to be responsible for synthesis of the outer-core (OC) component of the LOS (38). The OC locus (OCL) is located in the chromosome between the *aspS* and *ilvE* genes (16, 39). Each distinct gene cluster found between the flanking genes is assigned a unique number identifying the locus type (OCL1, OCL2, etc.), and to date, 14 different gene clusters (OCL1-12 (39) and OCL15-16 (40) have been identified. Nomenclature for OCL genes is also shown in Table 1, and Gtrs encoded at the OC locus are differentiated from KL-encoded Gtrs by the addition of OC to the name (GtrOC#). Generally, OC gene clusters fall into two broad families (Figure 1B), designated Group A and Group B, defined by the presence of *pda1* and *pda2* genes, respectively (39).

Several studies have highlighted the extremely plastic nature of the *A. baumannii* genome, revealing very poor correlation between KL and OCL types and other genomic features including sequence type (2, 16, 23, 41-43). Therefore, the most valuable framework for tracing important genetic lineages of *A. baumannii* currently involves a combination approach, including phylogenetic analysis with multi-locus sequence typing (MLST) using both Institut Pasteur and Oxford schemes, resistance and virulence gene mapping, and K and OC locus typing (2, 40-43). Bioinformatics tools and databases currently exist for MLST and resistance gene typing, allowing multiple genomes to be processed quickly. However, the lack of computational tools and databases to rapidly extract interpretable, actionable information about K- and OC- loci from large data sets is a current bottleneck.

Recently, a computational tool, named *Kaptive*, was developed to rapidly identify reference K and O loci in *Klebsiella pneumoniae* species complex genome sequences taking as input a curated database of reference sequences and a query genome assembly (44, 45). Though the computational tool can be used to type loci in any species, a complete and curated compendium of appropriate, species-specific KL, OL or OCL sequences is needed. In the case of *A. baumannii*, such databases are not currently available.

Here, we present curated databases of annotated reference sequences for *A. baumannii* K and OC loci that are compatible with *Kaptive*, enabling rapid typing of genomes for this clinically significant pathogen. We evaluate the accuracy of this approach by comparison of K and OC locus calls for >630 genomes typed previously using manual methods.

Additionally, we apply this approach to type >3300 *A. baumannii* genomes retrieved from the NCBI database, highlighting the extent of K and OC locus variability in the broader population and among clinically important clonal complexes, and confirming that the vast majority of genomes harbour loci matching those in our reference databases.

## Materials and Methods

### K and OC reference sequences

Nucleotide sequences for reference isolates carrying each KL and OCL type were downloaded from NCBI non-redundant or WGS databases (accession numbers are listed in Tables S1 and S2). Where possible, whole genome sequences were assessed for the presence of the *A. baumannii-*specific *oxaAb* gene (GenBank accession number CP010781.1, base positions 1753305 to 1754129) to confirm the sequences were obtained from an *A. baumannii* isolate. A GenBank format file (.gbk) for each distinct locus type was prepared. This file includes the nucleotide reference sequence for the locus without flanking sequence, the annotations of all coding sequences in the locus, and citation(s) for the annotations and/or polysaccharide structural data, if available.

### Curated Kaptive databases

The individual KL files were concatenated into a multi-record GenBank-format file to produce a data set containing annotated KL reference sequences. Likewise, the OCL files were compiled to generate a separate data set. Both reference databases were integrated with the *Kaptive-Web* platform (http://kaptive.holtlab.net/), which enables users to submit their genome sequence queries to a browser and receive the output in a visual format, as described in detail previously (45). The KL and OCL databases have also been made freely available for download from the *Kaptive* github repository (https://github.com/katholt/Kaptive) for use with the command-line version of *Kaptive* (44), or other tools.

### Genome sequence collections

*Acinetobacter* genome assemblies from our collection for which the KL and OCL types had been previously determined via manual or automated sequence inspection (2, 41); and unpublished data) were used to assess the level of typing accuracy that could be achieved through the use of our novel databases with *Kaptive*. Paired-end Illumina read data (described in (2, 41) and available under BioProject accession PRJEB2801) were *de novo* assembled using SPAdes v 3.13.1 (46) and optimised with Unicycler v 0.4.7 (47). High-quality genome assemblies (n = 719) with a maximum contig number of 300 and minimum assembly length 3.6 Mbp were included in the analysis (cut-offs determined empirically by manual inspection of the contig number and assembly length distributions, respectively). These assemblies were assessed for *oxaAb* presence using BLASTn (>95% nucleotide sequence identity and >90% combined coverage) to confirm the *A. baumannii* species assignment. Confirmed *A. baumannii* sequences (n = 642) were analysed using both KL and OCL reference databases with command-line *Kaptive* v 0.7 (44) with default parameters.

The same method was used to test databases against 3412 genome sequences available in the NCBI non-redundant and WGS databases as of February 2019. These genome assemblies were bulk downloaded from NCBI as a compressed .tar file for local analysis. Genomes lacking *oxaAb* were removed prior to typing but quality control (QC) analysis as described above was applied to this data set only after typing was complete.

### Interpretation of Kaptive output

The *Kaptive* output is described in detail elsewhere (45). Briefly, *Kaptive* uses a combination of BLASTn and tBLASTn searches to identify the best matching reference locus for each query genome and indicates a corresponding confidence level. The latter is dependent on the BLASTn coverage and identity for the full-length reference locus, the number of reference locus genes (expected genes) or other genes (unexpected genes) found within the locus region of the query genome (determined by tBLASTn, default coverage cut-off ≥90%, identity ≥80%), and whether the locus is found on a single or multiple assembly contigs. A ‘perfect’ confidence match indicates that the locus was found in the query genome on a single contig with 100% coverage and 100% nucleotide identity to the best-match reference locus. ‘Very high’ confidence matches are those for which the locus is present in the query genome in a single assembly contig with ≥99% coverage and ≥95% nucleotide sequence identity to the best-match reference locus, and no missing or unexpected genes within the locus. ‘High’ confidence matches are defined as those for which the locus was found on a single contig with ≥99% coverage to the best-match reference locus, ≤ 3 missing genes and no unexpected genes within the locus. ‘Good’ confidence matches indicate that the locus was found on a single contig or split across multiple assembly contigs with ≥95% coverage to the best-match locus, ≤ 3 missing genes and ≤ 1 unexpected gene within the locus. ‘Low’ confidence matches indicate that the locus was found on a single contig or split across multiple assembly contigs with ≥90% coverage to the best-match locus, ≤ 3 missing genes and ≤ 2 unexpected genes within the locus. A confidence level of ‘None’ indicates that the match does not meet the criteria for any other confidence level.

### Distribution of K and OC loci

For NCBI genome assemblies, sequence types (STs) were assigned with the mlst script (github.com/tseeman/mlst) using the Insitut Pasteur scheme for *A. baumannii* (abaumannii_2 scheme) available at https://pubmlst.org/bigsdb?db=pubmlst_abaumannii_pasteur_seqdef. KL and OCL variation were visualised for STs with ≥20 isolate representatives with ‘good’ or better confidence matches called by *Kaptive*.

## Results

### KL and OCL numbering and nomenclature

The development of curated databases for numbered and fully annotated *A. baumannii* K- and OC- loci relies on the consistent application of a standardised nomenclature and numbering system for these loci. Here, the system developed for transparent annotation of both the K and OC loci (16) has been used. As new KL and OCL types with additional gene families have been discovered since 2013, the gene nomenclature has been extended and is summarised in Table 1. For consistency, K loci that were originally published using other nomenclatures or typing systems have been re-annotated, and where possible the corresponding GenBank entries have been updated with the permission of the original authors (see Table S1).

In several cases, KL types that differ only by a small portion of the locus have been found e.g. (16, 48) and examples are shown in Figure 2. In cases where structures have been determined, the locus difference is associated with changes in the composition or structure of the CPS (26, 27, 29, 31, 35, 49-54) but some locus differences are now known to have no effect on CPS structure (24, 55). As all differences in genetic content are relevant in epidemiological studies, all K loci comprising a unique combination of genes were distinguished with a new KL number.

### The curated KL reference database

The annotations for 92 of 128 KL types are publicly available. Curated annotations have been deposited into GenBank for 78 KL types, three of which were submitted as third party annotations (TPA) (see Table S1). An additional 14 sequences were extracted from genomes in the WGS database (see Table S1). Sequences for the remaining 37 KL types are not currently available in the public domain.

Complete annotations for the 92 publicly available *A. baumannii* K locus reference sequences spanning the full length of each gene cluster (between *fkpA* and *lldP*) were therefore compiled into a KL reference database for use with *Kaptive*. Where the only available representative of a KL type included an insertion sequence (IS), we substituted the sequence with a manually generated version with the IS and target site duplication removed in order to include a KL that represents the presumptive ancestral, non-modified sequence as is required for accurate typing by *Kaptive* (44). This was the case for KL types KL27, KL44, KL82, KL87, KL93, KL114, and KL118 (Table S1).

### The curated OCL reference database

The annotations for 12 different OCL types have been described in the literature (39). A complete list is found in Supplementary Table S2. However, only six of them were available in GenBank. The remaining six OCL sequences were identified in the WGS database, and the WGS accession numbers are available in Table S1. Complete annotations for the 12 publicly available OCL spanning the full length of the gene clusters (between *ilvE* and *aspS*) were combined into a single OCL reference database for use with *Kaptive*.

### Compatibility of the KL and OCL databases with Kaptive

To confirm the compatibility of the KL and OCL databases for *Kaptive*-based typing, we created two query sequence sets comprising FASTA sequences of the reference KL and OCL, respectively. *Kaptive* was applied to each of these query sets, and was able to successfully identify the correct locus in all cases (Dataset 1).

### Comparison of Kaptive assignments with previous KL assignments

We assessed the accuracy of *Kaptive*-based KL typing using our curated KL database by application to a collection of 642 *A. baumannii* genome assemblies (see Dataset 2), which had been typed previously using BLASTn plus manual inspection (2, 41; and unpublished data). For these assemblies, the confidence levels called by *Kaptive* were: 176 (perfect), 385 (very high), 28 (high), 53 (good), 0 (low) and 0 (none) (Figure 3A; Dataset 2). Notably, 561 matches were assigned ‘perfect’ or ‘very high’ confidence calls, demonstrating that *Kaptive* could very confidently assign a KL type to the majority (87.4%) of the 642 genome assemblies provided.

**Figure 3.**
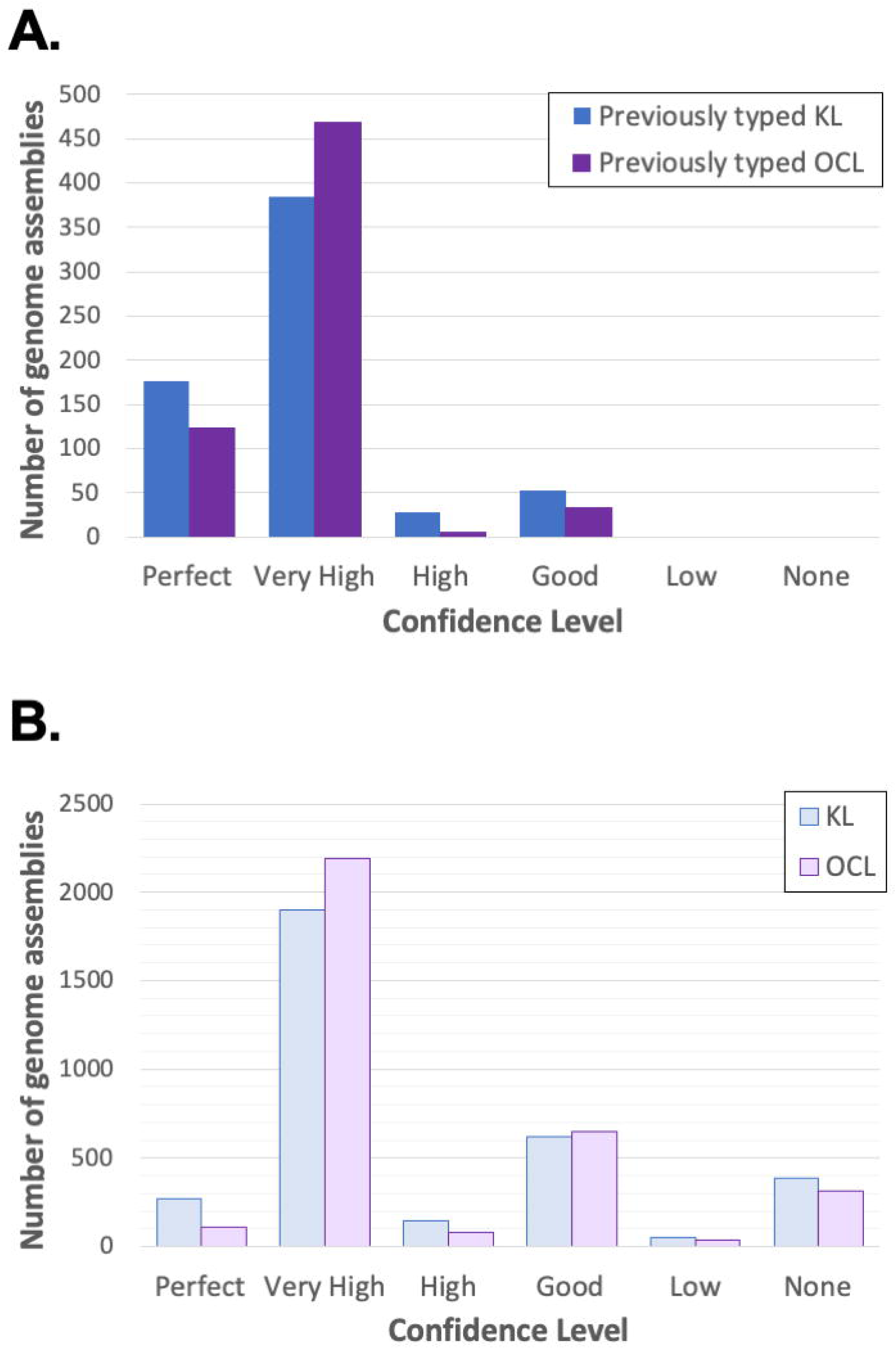
Breakdown of confidence levels for *Kaptive* locus calls using the *A. baumannii* KL and OCL databases. **A.** Results following database quality checking using private collection of 680 *A. baumannii* genome assemblies (Dataset 2). Colour key is shown in the top right corner. **B.** Results of applying the databases to 3412 genome assemblies available in NCBI databases (Dataset 3). Colour key is shown in the top right corner.

The 28 ‘high’ confidence matches each included one or more single base deletions within the locus leading to the interruption of a coding sequence, which *Kaptive* reports as one or more missing genes when the resulting tBLASTn matches have <90% coverage to the reference gene sequence. Such deletions may represent sequencing and/or assembly errors but may also represent true sequence variations with the potential to result in altered CPS structure. Since *Kaptive* is unable to distinguish these possibilities it reports the ‘missing’ gene and lowered confidence score in order to alert the user and facilitate further investigation.

Manual inspection of the relevant assembly graphs showed that 50 of 53 (94.4%) assignments with a ‘good’ confidence level were locus variants in which an IS had interrupted the KL gene cluster breaking it into two or more contigs in the query genome. The three remaining assemblies that were typed with a ‘good’ confidence level were also broken into multiple contigs that represented dead-ends in the assembly graphs, hence it was not possible to determine if these also represented IS variants or were simply the result of assembly problems e.g. due to low sequencing depth in the KL region of the genome.

Of the 642 assemblies with a KL type that was assigned previously, 641 (99.8%) were concordant and one (0.2%) was discrepant. The K locus of *A. baumannii* isolate BAL_266 had previously been described as KL63 (41). However, *Kaptive* assigned it to KL108 with a ‘very high’ confidence level (99.98% nucleotide sequence identity; 100% coverage). The sequence of this isolate was manually checked again and confirmed to be KL108. The KL63 and KL108 gene clusters are 97.96% identical across 95% of the locus, differing from each other only in ~1.3 kb segment in the central region that includes the *wzy* gene (Figure 2A).

This small difference between the two gene clusters was missed in the original manual typing but likely alters the linkage between the K units. This highlights the need to look for any regions of sequence difference when manually typing.

### Comparison of Kaptive assignments with previous OCL assignments

We also assessed the accuracy of OCL identification using our curated OCL database applied to the same collection of *A. baumannii* genomes. The OCL region of 631 of these had previously been typed using BLASTn plus manual inspection (2, 41; and unpublished data). The confidence levels for the OCL matches for the 631 typed genomes were: 124 (perfect), 469 (very high), 5 (high), 33 (good), 0 (low) and 0 (none) (Figure 3A; Dataset 2). As for the KL database, the large number of ‘perfect’ and ‘very high’ confidence matches (593, 94.0%) demonstrate the capacity of the OCL database to type the majority of genome assemblies provided as a query. Manual inspection confirmed that the five ‘high’ confidence matches included those with one or more base deletions in coding sequences, and the 33 ‘good’ matches represented variants of the corresponding reference sequences interrupted by one or more ISs. In this set, there were no discrepancies between the previous locus assignments and those determined by *Kaptive*.

### Application of KL and OCL databases for A. baumannii genome typing

As the KL and OCL regions in the majority of NCBI genome sequences have not yet been examined, the publicly available genomes provide a large dataset to begin to explore KL and OCL diversity in the species. Available genome assemblies of 3412 isolates annotated as *A. baumannii* in the NCBI non-redundant and WGS databases were first checked for the presence of the *oxaAb* gene to ensure correct assignment to the *baumannii* species. The *oxaAb* gene was absent from 34 assemblies (0.99%), and these were removed from the analysis bringing the total number of assemblies examined to 3378.

For the KL database, the confidence levels of the matches called by *Kaptive* were: 272 (perfect), 1901 (very high), 149 (high), 622 (good), 51 (low) and 383 (none). Among the 2944 genomes with KL confidence matches ‘good’ or better, there were 79 distinct KL types, 36 (45.6%) of which were identified in five or fewer genomes. Notably 13 of the loci included in the KL reference database were not identified among any of the genome assemblies retrieved from the NCBI database. The most common KL types were KL2 (713 of 2948 genomes, 24.2%), KL9 (343, 11.6%), KL22 (330, 11.2%), KL3 (294, 10.0%) and KL13 (155, 5.3%).

For the OCL database, the confidence levels were as follows: 108 (perfect), 2192 (very high), 80 (high), 645 (good), 39 (low) and 314 (none) (Figure 3B; Dataset 3). All 12 of the reference OC loci were identified among the 3029 genomes with OCL confidence matches ‘good’ or better. Among these genomes the most common OCL types were OCL1 (2086, 68.9%), OCL3 (272, 9.0%), OCL6 (157, 5.2%), OCL2 (150, 5.0%) and OCL5 (125, 4.1%).

Therefore, among the *A. baumannii* genomes retrieved from NCBI, KL and OCL calls were obtained for 87% and 90% of the assemblies, respectively. However, ‘low’ and ‘none’ confidence levels may result from poor quality sequence assembly and/or may indicate that a novel locus is present in the query assembly (44). Indeed, the application of the same quality control cutoff used for inclusion in our own data set (see above) revealed that 13/51 ‘low’ and 174/387 ‘none’ confidence matches for the KL assignments may be assemblies of poor quality. Similarly, 12/39 ‘low’ and 76/314 ‘none’ confidence matches for the OCL assignments did not meet the same quality control cutoff. Hence, it is recommended that users perform additional investigations to confirm the quality of their assemblies before excluding ‘low’ and/or ‘none’ confidence matches from their analyses.

### KL and OCL variation in clonal lineages

Variation in the KL and OCL in the major multi-drug resistant clonal lineages have largely been examined using small datasets (e.g. (2, 16, 38, 39)). For the GC2 lineage, these studies assessed diversity amongst isolates predominantly recovered from the same outbreak or region (41-43) or sporadic isolates (26, 56), limiting the ability to gain a complete picture of surface polysaccharide variation in this clone. Across these studies, at least 14 KL and 5 OCL have been reported in GC2. Of the 3386 genome assemblies we analysed here, 2016 belonged to ST2 in the Institut Pasteur scheme, representing the most common ST in GC2 and the largest group of isolates belonging to a single ST (6). Among the 2016 ST2 genomes, *Kaptive* identified 30 KL and 3 OCL (Figure 4) in those with confidence matches ‘good’ or better. The most common KL arrangements were KL2 (32.2%) and KL22 (14.4%), whereas OCL1 represented the most predominant OCL type (78.6%). Only one KL, KL63, was found in a single ST2 genome. For the remaining assemblies, 107 (5.3%) and 256 (12.7%) were assigned ‘low’ and ‘none’ confidence matches against the KL and OCL databases, respectively. These assemblies may be of poor quality or they may carry novel types but this was not further investigated.

**Figure 4.**
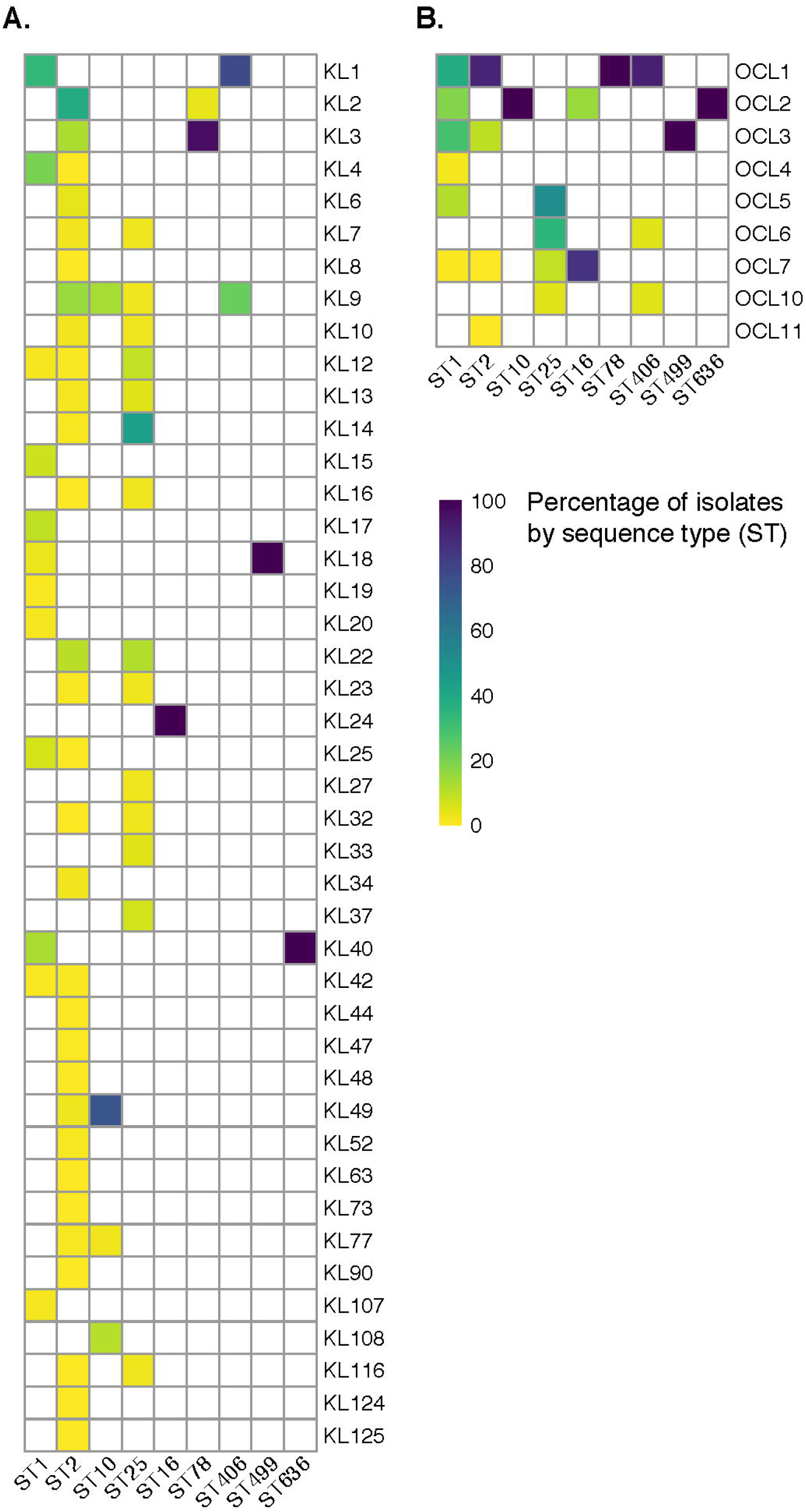
Distribution of K and OC loci by sequence type. Heat maps show the distribution of distinct K (**A**) and OC (**B**) loci among genomes assigned to nine common multi-locus sequence types (STs). Coloured shading indicates the percentage of isolates belonging to a given ST that were assigned a given K or OC locus type, as indicated by the colour legend*. A. baumannii* genome assemblies were retrieved from the NCBI database; only confirmed *A. baumannii* for which both K and OC loci were assigned by *Kaptive* with confidence level “Good” or better are shown (n = 2002; 125 ST1, 1669 ST2, 46 ST10, 20 ST16, 43 ST25, 28 ST78, 22 ST406, 29 ST499, 20 ST636).

KL and OCL diversity have also previously been reported for the other major clonal lineage, GC1. An in-depth study of 45 *A. baumannii* GC1 isolates identified 8 KL and 5 OCL types in this clone (2), with one additional KL type found in a subsequent study (57). In the set of 3386 genome assemblies, we found 134 that belong to ST1, which represents the most common GC1 sequence type. *Kaptive* identified a total of 10 KL and 6 OCL types in the ST1 lineage (Figure 4), expanding the number of distinct types observed previously. Among these ST1 genomes, the most common KL types were KL1 (31.3%) and KL4 (18.7%), while the most common OCL were OCL1 (36.6%), OCL2 (17.2%) and OCL3 (29.1%). KL19 and KL42 and also OCL7 were found in single isolates.

We also examined a further seven STs for which there were ≥20 isolate representatives with confident *Kaptive* matches (‘good’ or better). Of these STs, ST10 included the largest number of genome assemblies (47 of 3386 assemblies), and 4 KL and 1 OCL type were found in this group. ST25, the second largest group with 46 assemblies had very high variation with 14 KL and 4 OCL types. ST406 (22 assemblies) also included 4 OCL types but only 2 KL. However, one of two KL types and one or two OCL types were found in ST16 (20 assemblies), ST78 (29), ST499 (29) and ST636 (20). Notably, specific KL and OCL types were not confined to single STs, with several locus types found in more than one ST.

## Discussion

In this study, we present *Kaptive* compatible databases of annotated reference sequences for *A. baumannii* K and OC loci, extending the utility of *Kaptive* and broadening the ability of researchers, clinicians and public health professionals to analyse genome data sets. Using these databases, *Kaptive* was able to confidently and accurately assign KL and OCL types to the majority of *A. baumannii* genome assemblies examined. Among >630 *A. baumannii* genomes typed previously using manual methods, only a single discrepancy between the previous KL assignment and that of *Kaptive* was identified. This was traced to an error in the previous manual assignment which had overlooked a small genetic replacement within the locus. As sequence replacements of < 2 kb are common in *A. baumannii* KL and OCL regions (examples shown in Figure 2), the ability of *Kaptive* to correctly identify the KL type demonstrated the stringent nature of the tool and the quality of the databases described herein. The KL and OCL databases were also used to probe the collection of *A. baumannii* genome assemblies available through NCBI GenBank and WGS databases. *Kaptive* was able to confidently assign locus types to more than 87% of these genome assemblies, indicating that the databases capture the majority of common KL and OCL types. However, to confirm the locus calls, all *Kaptive* assignments should be checked for length discrepancies that would reveal missing expected genes, and/or the presence of additional genes or IS in the locus.

The remaining genomes that could not be confidently assigned a locus type (13% KL and 10% OCL unassigned) may include genomes with low coverage and/or poor assembly quality in the KL and/or OCL genome regions. Alternatively, these genomes may carry loci that are not represented in the current reference databases. In these cases, users are encouraged to undertake further investigations e.g. by manual inspection of the assembly and/or assembly graphs and comparisons to the best-matching reference loci using visualisation tools such as Artemis Comparison Tool (58) and Bandage (59). Further work will be needed to identify and include further novel loci and the databases will be continuously updated as sequences and annotations for further KL and OCL types become available. We encourage users to contact us via the *Kaptive-Web* website and/or the *Kaptive* github page to submit novel loci for the assignment of KL and OCL numbers and addition to the publicly available databases.

The typing system and the databases have been designed strictly for use in *A. baumannii* and therefore users are encouraged to check the origin of their sequences to ensure reliable results. The presence of the intrinsic *oxaAb* gene in the genome sequence can be applied as a simple check to confirm a sequence is from an *A. baumannii* isolate prior to use of the databases, bearing in mind that it may be missing from poor quality assemblies. However, this does not preclude the use of the *A. baumannii* KL and OCL databases on other species of *Acinetobacter*. Though not all locus types found in other species will be represented in the databases, K or OC loci with high similarity to those found among *A. baumannii* can be easily identified (see examples in Dataset 3). Hence, the *A. baumannii* databases may assist identification and annotation of the specific genetic content of loci in other *Acinetobacter* species.

It should be noted that the KL does not predict the structure of the CPS, though it does include information about the possible number and identity of sugars present. The CPS structure for each KL must be determined directly as in a number of cases additional genes involved in capsule synthesis are found outside the locus (28, 51, 54). Hence the KL type is only a starting point for predicting if a particular isolate might be susceptible to a particular phage. However, the potential power of KL and OCL typing as epidemiological tools is highlighted by the analysis of KL and OCL found in single STs. KL and OCL typing have previously proven valuable in dissecting the evolution of two major global clones (2, 41-43). However, in most studies the genomes were typed using a time intensive manual process, which imposed a considerable limitation on the scale of datasets that could be explored. In contrast, in this work we were able to use the automated method implemented in *Kaptive* to type the K and OC loci of 1000s of genomes, including 134 GC1 and 2016 GC2 revealing even more extensive variation, which is likely to be driven by exchange of locus sequences via recombination in both clones. Given that the available genomes are drawn from a biased, convenience sample of genomes deposited in NCBI (6), they still may not reflect the true variation in these clones. Similar high levels of variation were found in two other clones (ST10 and ST25), suggesting that they are subject to similar molecular evolutionary processes. In contrast, there appeared to be limited KL and OCL variation among ST16, ST78, ST406, ST499 and ST636.

The findings reported here clearly demonstrate the utility of our novel KL and OCL databases to facilitate rapid and accurate typing of *A. baumannii* surface polysaccharide synthesis loci. This information can be used to distinguish lineages within the global clonal complexes (2, 41, 57) and hence provide valuable information for epidemiological studies, as well as essential information to guide the design of novel treatment or control strategies targeting *A. baumannii* capsules and lipooligosaccharides.

