## Supplementary material for "Identification of *Acinetobacter baumannii* loci for capsular polysaccharide (KL) and lipooligosaccharide outer core (OCL) synthesis in genome assemblies using curated reference databases compatible with Kaptive": Table S1

**Table S1.** GenBank and WGS accession numbers for the sequences used to construct the *A. baumannii* KL database

| **K locus** | **Reference isolate** | **GenBank or WGS accession number** | **Base range** | **Reference** |
| --- | --- | --- | --- | --- |
| KL1 | A1 | CP010781.1 | 87963..110544 | (1, 2) |
| KL2 | A74 | KJ459911.1 | 3057..27820 | (1, 3, 4) |
| KL3 | D86 | KF793926.1 | 3248..25749 | (1) |
| KL4 | D78 | JN409449.3 | 3247..31285 | (1, 2, 5) |
| KL5 | SDF | BK010760 ^2^ | 1..25029 | (1, 6, 7, 8, 9) |
| KL6 | F4 | KF130871.1 | 3248..26268 | (1, 4, 10) |
| KL7 | SGH0701 | KX011025.2 | 3058..28708 | (1, 6, 8) |
| KL8 | BAL097 | KX712116.2 | 724..32274 | (1, 6, 11) |
| KL9 | RUH134 | JN247441.4 | 3250..28148 | (1) |
| KL10 | BAL_030 | KY434633.1 | 3058..28938 | (1, 12) |
| KL11 | J9 | KF002790.2 | 745..26607 | (13) |
| KL12 | D36 | JN107991.2 | 3247..38792 | (2, 6, 14) |
| KL13 | Ab689 | MF522810.1 | 724..36552 | (6, 15) |
| KL14 | D46 | KF030679.2 | 724..21932 | (16, 36) |
| KL15 | A85 | KC118540.6 | 9179..37114 | (2, 17, 18) |
| KL16 | D4 | MF522813.1 | 724..23467 | (19) |
| KL17 | G7 | KC118541.2 | 3247..25474 | (2) |
| KL18 | Ab762 | MF522811.2 | 724..24288 | (20) |
| KL19 | RBH2 | KU165787.1 | 724..22129 | (21, 22) |
| KL20 | A388 | JQ684178.2 | 3250..32301 | (2, 23) |
| KL21 | G21 | MG231275.1 | 3058..33061 | (23) |
| KL22 | LUH5537 | KC526920.2 ^3^ | 3248..27716 | (24) |
| KL23 | Ab836 | MF522812.1 | 724..25107 | (4) |
| KL24 | RCH51 | KX756650.1 | 724..25150 | (12) |
| KL25 | AB5075 | BK008886 ^2^ | 1..23498 | (2, 25) |
| KL26 | Ab902 | MF522809.1 | 724..28488 | (13) |
| KL27^1^ | 4190 | KT266827.1 | 382..31000 | (6, 26) |
| KL28 | Ab908 | MF522807.1 | 724..26378 | (6) |
| KL29 | 2007-16-27-01 | AMIR01000046.1 ^4^ | 11241..36611 | (13) |
| KL30 | NIPH 190 | MN166189.1 | 1..20785 | (27) |
| KL31 | OIFC0162 | AMFH01000019.1 ^4^ | 107741..131132 | (4) |
| KL32 | BAL 058 | KT359615.1 | 724..22514 | (3) |
| KL33 | NIPH 67 | MN166195.1 | 1..22775 | (4, 28) |
| KL34 | BJAB07104 | CP003846.1 ^4^ | 94048..118441 | (23) |
| KL35 | LUH5535 | KC526896.2 ^3^ | 4373..27270 | (18) |
| KL36 | Naval-72 | AMFI01000021.1 ^4^ | 38074..64065 | (13) |
| KL37 | UV1036 | KX712115.1 | 724..21811 | (24, 36) |
| KL38 | 4300STDY7045887 | UFOG01000008.1 ^4^ | 32622..55633 | - |
| KL39 | AB_2008-15-71 | AMHP01000019.1 ^4^ | 57921..78767 | (21) |
| KL40 | D141c | KP100029.1 | 724..24731 | (2) |
| KL41 | Naval-57 | AMFP01000100 ^4^ | 42104..65411 | - |
| KL42 | NIPH24 | MN166194.1 | 1..22901 | (4, 29) |
| KL43 | NIPH 60 | MN166192.1 | 1..19982 | (30) |
| KL44^1^ | NIPH 70 | MN148385.1 | 1..26811 | (6, 26) |
| KL45 | NIPH 201 | MN166190.1 | 1..21078 | (27) |
| KL46 | NIPH 329 | MK609549.1 | 1..23617 | (4, 9) |
| KL47 | UV1043 | KX661320.1 | 724..22352 | (30) |
| KL48 | NIPH 615 | MN166191.1 | 1..20705 | (27) |
| KL49 | BAL_173 | KT359616.1 | 724..32883 | (3, 6) |
| KL50 | OIFC047 | AMFW01000039.1 ^4^ | 35120..58660 | - |
| KL51 | WM98c | MN148384.1 | 1..22098 | - |
| KL52 | H32 | KY434632.1 | 3057..23978 | - |
| KL53 | D23 | MH190222.1 | 736..21707 | (22) |
| KL54 | RCH52 | MG867726.1 | 724..32088 | (11) |
| KL55 | BAL 204 | MN148381.1 | 1..24890 | - |
| KL57 | BAL_212 | KY434631.1 | 724..26129 | (31) |
| KL58 | BAL_114 | KT359617.1 | 724..25347 | (3) |
| KL60 | BAL_329 | MN148382.1 | 1..24516 | (12) |
| KL63 | BAL_103 | KX712117.2 | 724..32678 | (6) |
| KL73 | SGH0703 | MF362178.1 | 724..36545 | (15, 32) |
| KL74 | BAL 309 | MN148383.1 | 1..25336 | - |
| KL77 | MSHR_188 | MK370019.1 | 1..23882 | (20) |
| KL80 | LUH3712 | KC526914.2 ^3^ | 3090..27616 | (12) |
| KL81 | LUH3713 | KC526916.2 ^3^ | 3247..29414 | (10) |
| KL82^1^ | LUH5534 | KC526908.2 ^3^ | 3248..23847 | (33) |
| KL83 | LUH5538 | KC526898.2 ^3^ | 538..26632 | (13) |
| KL84 | LUH5540 | KC526902.2 ^3^ | 3252..26477 | - |
| KL85 | LUH5543 | KC526913.2 ^3^ | 3089..27646 | - |
| KL87^1^ | LUH5547 | KC526918.2 ^3^ | 3249..32444 | (13) |
| KL88 | LUH5548 | KC526910.2 ^3^ | 3193..23919 | (30) |
| KL89 | LUH5552 | KC526919.2 ^3^ | 3159..27654 | (12) |
| KL90 | LUH5553 | KC526917.2 ^3^ | 3191..26854 | (8) |
| KL91 | 1053 | KM402814.1 | 1832..24537 | (34) |
| KL93^1^ | B11911 | BK010902 ^2^ | 1..27476 | (35) |
| KL102 | MSHR_200 | MK370021.1 | 1..21075 | (20) |
| KL105 | 625974 | JEXD01000015.1 ^4^ | 32795..58839 | (13) |
| KL106 | 219_ABAU | JVPN01000008.1 ^4^ | 31656..58916 | (13) |
| KL107 | MSHR_183 | MK370022.1 | 1..22790 | (20) |
| KL108 | MSHR_204 | MK370023.1 | 1..31284 | (20) |
| KL109 | MSHR_192 | MK370024.1 | 1..24118 | (20) |
| KL110 | MSHR_203 | MK370025.1 | 1..34861 | (20) |
| KL111 | MSHR_53 | MK370026.1 | 1..23163 | (20) |
| KL112 | MSHR_54 | MK370027.1 | 1..25441 | (20) |
| KL113 | MSHR_8 | MK370028.1 | 1..29261 | (20) |
| KL114^1^ | MSHR_89 | MK388214.1 | 1..27862 | (20) |
| KL116 | MAR-303 | MK339425.1 | 3275..23797 | (36, 37) |
| KL118^1^ | TG00314 | ASER01000021.1 ^4^ | 37142..66310 | (23) |
| KL119 | ARLG1794 | NGGP01000084.1 ^4^ | 30036..54769 | (31) |
| KL120 | ABBL011 | LLCR01000062.1 ^4^ | 15005..38992 | (9) |
| KL124 | ABUH511 | NCXX01000026.1 ^4^ | 32588..54524 | (33) |
| KL125 | MAR13-1452 | MH306195.1 | 5072..32981 | (38) |
| KL128 | KZ-1093 | MK339428.1 | 3274..5469 | (37) |

^1^ Sequence has been modified to remove IS or IS remnants for inclusion in the KL database

^2^ Third Party Accession (TPA) including current nomenclature and naming

^3^ Old annotations updated in this study consistent with established nomenclature scheme (1).

^4^ GenBank or WGS sequence not including current nomenclature and naming
