## Supplementary material for "Identification of *Acinetobacter baumannii* loci for capsular polysaccharide (KL) and lipooligosaccharide outer core (OCL) synthesis in genome assemblies using curated reference databases compatible with Kaptive": Table S2

**Table S2.** GenBank and WGS accession numbers for the sequences used to construct the *A. baumannii* OCL database

| **OC locus** | **Reference isolate** | **GenBank or WGS accession number** | **Base range** | **Reference** |
| --- | --- | --- | --- | --- |
| OCL1 | A1 | CP010781.1 | 3366405..3375181 | (1-3) |
| OCL2 | D36 | CP012952.1 | 646907..655270 | (1, 3) |
| OCL3 | A85 | CP021782.1 | 3464825..3473737 | (1, 3) |
| OCL4 | A388 | CP024418.1 | 642040..650673 | (3) |
| OCL5 | G21 | MG231275.1 | 37120..46166 | (3) |
| OCL6 | D46 | KF030679.1 | 28675..37977 | (3) |
| OCL7 | OIFC035 | AMTB01000038.1 ^1^ | 221522..230586 | (3) |
| OCL8 | OIFC111 | AMFY01000013.1 ^1^ | 222496..228777 | (3) |
| OCL9 | Naval-72 | AMFI01000027.1 ^1^ | 34336..40843 | (3) |
| OCL10 | AB_TG27343 | AMIS01000032.1 ^1^ | 97841..111169 | (3) |
| OCL11 | TG22204 | ASFV01000009.1 ^1^ | 33588..43841 | (3) |
| OCL12 | NIPH 410 | ATGJ01000006.1 ^1^ | 47175..57655 | (3) |

^1^ GenBank or WGS sequence not including current nomenclature and naming
